# Novel Laser Technology Enables 10x Faster SRS Imaging and Rapid Tuning in Biological Samples

**DOI:** 10.1101/2025.01.31.635524

**Authors:** Lenny Reinkensmeier, René Siegmund, Mark Bates, Gero Stibenz, Peter Trabs, Stefan Popien, Ingo Rimke, Sandro Heuke, Alexander Egner

## Abstract

Stimulated Raman Scattering (SRS) microscopy was developed for the label-free detection of molecular groups, addressing the speed limitations of spontaneous Raman microscopy. Standard SRS microscopy typically operates with laser sources at an 80 MHz repetition rate and a color-tuning speed of approximately 0.1 Hz to target different molecular groups. Here, we present a novel laser system that overcomes these speed limitations, achieving an order-of-magnitude improvement in both color-tuning and imaging speed. Our system features a reduced repetition rate of 40 MHz, enabling SRS imaging that is ten times faster than standard systems while maintaining the same average power at the sample. This is achieved through increased pulse energy and laser modulation at half the repetition rate. Furthermore, the system provides nearly ten times faster color-tuning across an extended range (660–1010 nm) by employing angle-tuning of nonlinear crystals instead of temperature-tuning. The improved performance is demonstrated in direct comparison with a standard SRS laser system, showcasing the potential for significantly enhanced imaging capabilities.

## 1. Introduction

Coherent Raman Scattering (CRS) microscopy has emerged as a transformative tool for label-free, high-resolution chemical imaging, enabling the rapid visualization of molecular structures without the need for external labels [1, 2]. By exploiting nonlinear optical processes, CRS techniques probe molecular vibrations to provide intrinsic chemical contrast, making them invaluable in diverse fields ranging from materials science to biology [2, 3]. Among CRS modalities, Stimulated Raman Scattering (SRS) microscopy has gained particular prominence due to its linear dependence on molecular concentration, absence of non-resonant background signals, and high sensitivity, which together enable quantitative imaging of complex biological and chemical specimens [4].

SRS microscopy relies on the interaction of two synchronized laser beams, known as the pump and Stokes beams. When the energy difference between these beams matches the vibrational frequency of a specific molecular bond, coherent Raman signals are created at the wavelength of the incident lasers and are further amplified by interference with these lasers [5, 6]. This process, referred to as stimulated Raman loss (SRL) or stimulated Raman gain (SRG), increases sensitivity by orders of magnitude compared to spontaneous Raman scattering, enabling the detection of weak molecular vibrations with unparalleled speed and precision [4, 7]. Fast switching one laser on and off allows for detecting the SRS signal as a small (10^−4^ − 10^−6^) power modulation superimposed on the other laser. The detection of these subtle power variations requires laser sources with a flat beam profile (good M^2^ value), high temporal and spatial stability, and laser excess noise below the shot-noise at power levels above 10 mW to achieve high signal-to-noise ratios [8].

The performance of SRS microscopy is inherently tied to the properties of the employed laser systems. Critical parameters, such as pulse energy, spectral resolution, tuning speed, and wavelength coverage, directly affect imaging capabilities [9,10]. Broader spectral coverage allows access to both high-frequency vibrational modes, such as CH- and OH-stretching vibrations, and the fingerprint region, which contains unique molecular signatures [11]. Additionally, rapid wavelength tuning is essential for hyperspectral imaging, enabling the investigation of biochemical heterogeneity in tissues and other complex samples [3, 12].

Advancements in laser technology have played a pivotal role in the evolution of SRS microscopy. Historically, light sources for SRS faced major constraints, requiring at least one shot-noise limited laser output, which imposed strict limitations on laser cavity design, pulse-picking technologies, and amplification. While many manufacturers focused on reducing laser noise, few addressed the critical parameter of repetition rate, despite its inverse quadratic impact on SRS imaging speed. Recent innovations, including the reduction of repetition rates and increased pulse energies, have fundamentally shifted this paradigm [2, 10]. These developments have facilitated applications in live-cell imaging, tissue analysis, and materials characterization, cementing SRS as a cornerstone of modern optical microscopy [4, 6].

In this study, we present a comparative analysis of two advanced laser systems for SRS microscopy: a commercially established benchmark system and a newly developed prototype. The prototype features innovations such as reduced repetition rates, higher pulse energy, and a redesigned OPO cavity, aiming to improve SNR and accelerate wavelength tuning. These enhancements are designed to address common limitations in throughput and spectral imaging flexibility while maintaining high stability and spectral resolution [13]. Through quantitative measurements and qualitative imaging experiments, we evaluate the impact of these advancements on SRS performance. Biological samples, including onion epidermal cells and oyster prismatic layers, serve as test cases to demonstrate the system’s ability to resolve fine structural details and enable rapid sequential imaging across multiple Raman bands [14, 15]. This work underscores the transformative potential of technological innovation in expanding the capabilities of SRS microscopy for high-sensitivity, high-throughput chemical and biological imaging.

## 2. Theory

Stimulated Raman Scattering (SRS) is a nonlinear optical process whose efficiency depends not only on the sample’s intrinsic properties but also strongly on the spatial and temporal intensity distributions of the pump and Stokes beams. In contrast to spontaneous Raman scattering, which is inherently weak, SRS benefits from the coherent interaction of two synchronized laser pulses (pump and Stokes). By tuning the energy difference between these pulses to match a specific molecular vibration, the Raman response is amplified by orders of magnitude. This process transfers net energy from the pump to the Stokes beam, thus boosting the otherwise weak Raman signal well above the detection threshold. SRS is often called a two-photon process, although it differs conceptually from other two-photon phenomena such as two-photon fluorescence (TPF). In TPF, two photons are absorbed (nearly) simultaneously to reach an electronic excited state, whereas in SRS, one pump photon is effectively converted into a Stokes photon, providing a resonant vibrational excitation of the sample. This distinction underlines why SRS is often termed a “vibrationally resonant” process, since the difference frequency between pump and Stokes beams matches a specific molecular vibration. To quantitatively compare signal-to-noise ratios from different laser sources, we must ensure that any relevant differences are well-characterized or do not significantly affect the measurements. Additionally, understanding the various noise contributions is essential for accurate data interpretation. Although fundamental SRS equations are well established, we provide a concise summary to lay the foundation for our analysis. In this study, we focus on stimulated Raman loss (SRL), although the derived results also apply to stimulated Raman gain (SRG).

### 2.1 Pulse energy Loss in SRS

Gao et al. [13] have shown that the stimulated Raman scattering rate (in photons per second) for a specific vibrational mode, exposed to pump and Stokes intensities *I*_p_ and *I*_s_, can be expressed as

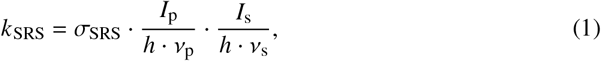

where σ_SRS_ is the molecule’s absolute stimulated Raman cross-section (with units of GM, where 1 GM = 10^−50^ cm^4^ ·*s*· photon^−1^), *h* is Planck’s constant, and *v* _*p*_ and *v*_*s*_ denote the respective light frequencies of the pump and Stokes beams.

This interaction leads to a decrease in the pump beam’s power, quantified by:

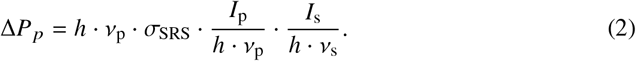

To determine the contribution of all molecules within the focal volume to the power loss of the pump beam, we require the intensity distribution in the focus of an objective lens [16, 17]:

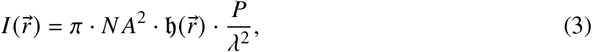

where *N A* = *n* sin α is the numerical aperture of the objective lens, 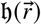 is the point spread function (PSF) normalized to 1 at the geometric focus, *P* is the power coupled into the back aperture of the lens and *λ* is the wavelength.

Because the relative change in pump and Stokes power is typically on the order of 10^−4^ to 10^−6^ [4], we can neglect their impact on the PSF in most practical scenarios, a concept referred to as the small-depletion assumption. Consequently, the overall power loss within the focal volume can be written as:

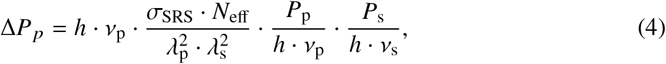

where we define *N*_eff_ as the effective number of Raman-active molecules within the focal volume. As discussed in [18, p. 111], for samples with a uniform concentration of scatterers, the SRS signal—and therefore *N*_eff_—is independent of the NA. However, for very high NA values, where the paraxial approximation no longer fully applies, it has been experimentally shown for two-photon excitation that the nonlinear excitation potential decreases with increasing NA [19] due to changes in the focal intensity distribution. Furthermore, we note that at very high laser intensities or under strongly resonant conditions, the small-depletion assumption may no longer hold.

Because SRS is a nonlinear Process, short-pulse lasers with pulse durations around one picosecond are commonly used in SRS microscopy. This shifts emphasis from pump power loss to the more significant pulse energy loss as the beam passes through the focal volume. The time-dependent power of a single pulse with energy *E* is expressed as:

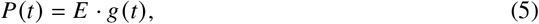

where *g* (*t*) , with units of *s*^−1^, characterizes the temporal shape of the pulse. We normalize *g* (*t*) so that its integral equals one, ensuring the pulse energy is *E*. The total energy loss of the pump pulse due to SRS within the focal volume is thus given by:

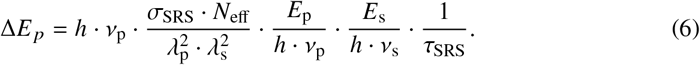

where we define the effective SRS pulse length, τ_SRS_, as:

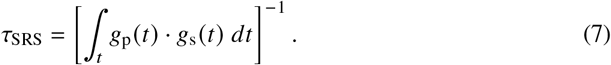

For ideal, temporally overlapping rectangular pump and Stokes pulses of length τ, the effective SRS pulse length is equal to τ. For two Gaussian pulses of lengths τ_p_ and τ_s_, the effective SRS pulse length at optimal overlap is 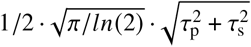.Thus, τ_SRS_ corresponds to the pulse length of ideal rectangular pump and Stoke pulses that would generate the same SRS signal.

### 2.2 Lock-in detection: Signal and noise in SRS

SRS microscopy operates at high photon flux, relying on the detection of subtle energy variations within an intense laser beam. Under standard conditions, the pump beam’s relative energy loss is on the order of 10^−4^–10^−6^ [4]. Such small variations are often obscured by laser and electronic noise, making direct detection difficult. To improve detection sensitivity, the excitation laser is modulated at several megahertz, and the SRS signal is demodulated at the same frequency. This approach moves the detection away from the electronic noise range (1/ *f* noise) and substantially minimizes laser excess noise.

In a typical SRS setup, pump and Stokes beams are accurately synchronized at a repetition rate *f*_rep_. For Stimulated Raman Loss (SRL) applications, the Stokes beam undergoes additional modulation at a frequency of *f*_mod_. This modulation is then transferred to the pump beam according to Eq. 6, facilitating its detection by the combination of a photodetector and a lock-in amplifier. Following Audier et al. [8], the lock-in output voltage is:

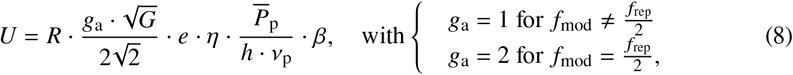

where *R* is the load resistor of the lock-in amplifier (input resistance of the ADC), *G* is the power gain of the amplifier, *e* is the elementary charge, and *η* is the quantum efficiency of the detector, 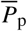 is the average power of the pump beam and *β* is the relative SRS loss Δ*E*_p_ / *E*_p_. The factor of two for *g*_a_ arises from optimized signal amplification during lock-in detection [20]. The lock-in amplifier multiplies the time trace of the pump power with a reference signal, typically a sine wave. When *f*_mod_ = *f* _rep_ ^2^, the pump pulse signal is consistently multiplied by +1 or 1, resulting in the maximum SRS signal contrast for the Stokes laser being switched on or off. If *f*_mod_ = *f*_rep_/ 2 is not satisfied, the multiplication with the sine wave reference results in factors varying across the full range from − 1 to +1. This includes cases where the Stokes laser is partially switched on but zero amplified, yielding an average multiplication factor of ±1/2.

Using Δ*E* _*p*_ from Eq. 6 and dividing it by *E* _*p*_ yields *β*. Substituting *β* into Eq. 8 and using the relationship 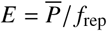then allows us to express *U* in terms of the average powers 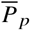 and 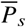,as shown below:

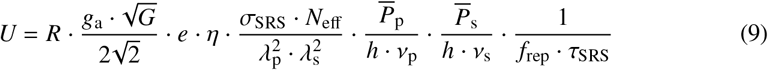

Notably, *U* is proportional to *N*_eff_, making it a common choice for representing SRS images.

The output signal of the amplifier is naturally subject to noise. Following Audier et al. [8], the variance of the output voltage is given by:

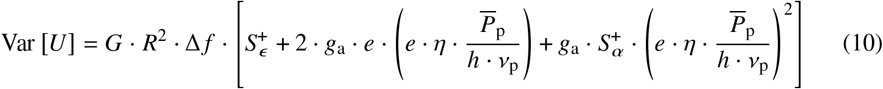

Consequently, the noise consists of three components: electronic noise, shot noise, and excess (classical) laser noise. Here, 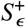 represents the absolute variance of the electronic noise as a function of the current at *f*_mod_, while 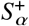 denotes the relative variance of the laser’s classical noise, expressed in terms of power, also at *f*_mod_. The term Δ *f* is the detection bandwidth of the lock-in amplifier or, with further averaging of the digitized output signal, the inverse of the pixel dwell time (Δ *f* = 1 / τ_pdt_). Note that when *f*_mod_ = *f*_rep_ / 2, the shot noise contribution encounters the same amplification as the lock-in output voltage, reflecting the increased modulation depth. Equation 10 specifically implies that the noise in SRS microscopy is purely dependent on the characteristics of the employed electronics (including the photodetector) and the laser used, and not on the SRS signal itself.

If the electronics’ noise is sufficiently low and the laser’s excess noise is below shot noise, then the noise floor is determined solely by shot noise, allowing the contributions of electronic and excess noise to be neglected. In the context of SRS, the signal-to-noise ratio (SNR) can be defined in different ways depending on the focus of the analysis: as SNR_std_ = *U*/ std [*U*] , which scales with concentration, or as SNR_var_ = *U*^2^ /Var [*U*] , which scales with time. Since this paper emphasizes speed, we reference the SNR using SNR_var_:

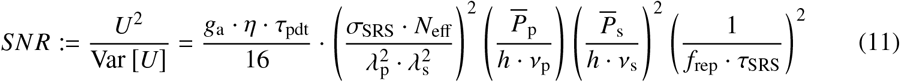

Hence, the SNR in SRS increases with higher average powers of both pump and Stokes beams, but decreases with increasing repetition rate *f*_rep_ and effective SRS pulse length τ_SRS_. In practice, balancing these parameters is crucial for optimizing the detection limit without compromising sample integrity. For instance, lowering *f*_rep_ at constant average power raises the pulse energy and thus the SRS signal, while tuning τ_SRS_ helps match the Raman linewidth for optimal spectral resolution. Moreover, adjusting the τ_pdt_ also affects the achievable SNR, since a longer pixel dwell time lowers the noise floor but reduces imaging speed.

## 3. Material and methods

### 3.1 Laser sources

SRS microscopy requires two laser pulses at different wavelengths with an energy difference matching the vibrational band of interest. These pulses must have precise temporal and spatial overlap at the sample. State-of-the-art laser sources utilize short picosecond pulses with bandwidths around 10 cm^−1^ to match typical Raman line widths, ensuring optimal excitation and spectral resolution. Operating in the near-infrared (NIR) range—the first optical window—allows for significant penetration depth and minimal scattering. By employing a Stokes wavelength around 1 µm in the SRL regime, where the SRS pump is detected, one can effectively use large-area, low-noise silicon photodiodes at their peak spectral sensitivity.

Jitter-free pump and Stokes pulses are essential for achieving broad tunability across the entire fingerprint region and high-frequency Raman modes, such as CH_*x*_ or OH vibrational bands ranging from below 500 to above 3500 cm^−1^. This is accomplished using synchronously pumped optical parametric oscillators (OPOs), which can be either solid-state or fiber-based. Solid-state OPOs offer advantages over fiber-based four-wave mixing OPOs, particularly in achieving inherently low noise levels—down to the shot noise limit in the MHz range—due to their higher intracavity power and noise averaging over multiple round trips in air cavities. This low-noise performance is further enhanced at high modulation and detection frequencies (e.g. 20 MHz), enabling video-rate SRS imaging.

Both light sources in this study are solid-state OPOs, pumped by frequency-doubled, mode-locked picosecond Yb lasers to generate the SRS pump beam. A part of the fundamental beam of the Yb-laser is modulated at 20 MHz to provide the Stokes beam. These sources feature automatic wavelength tuning, power attenuation, stabilization, and integrated temporal and spatial overlap of the pump and Stokes beams, enabling straightforward integration into a microscope.

The reference light source used is the picoEmerald S (APE, Germany), which employs a tuning mechanism based on temperature-tuned noncritical phase matching, as described in [21]. For small wavelength adjustments, tuning takes only a few seconds; however, larger wavelength changes can take up to 2 minutes due to the temperature tuning process. Other key parameters include 2 ps pulse lengths and an 80 MHz repetition rate.

To enhance the sensitivity of SRS and significantly accelerate tuning speed, we developed a prototype laser called picoEmerald FT. While maintaining a pulse duration in the range of 2 ps, the repetition rate is reduced from 80 MHz to 40 MHz. By maintaining the same average laser power, this change doubles the pulse energy, thereby enhancing the SRS signal. Since the modulation frequency of 20 MHz is phase-locked to the 40 MHz repetition rate, this synchronization, along with the increased pulse energy, leads to a theoretical increase in the SNR by a factor of 8 (see Eq. 11). Additionally, the OPO’s tuning mechanism is modified from temperature-based to mechanical tuning of the nonlinear crystal.

### 3.2 Microscope setup

The two SRS-laser sources, picoEmerald S and picoEmerald FT, were coupled into a custom built laser scanning microscope consisting of a beam scanner (Quad Scanner, Abberior Instruments, Germany) and an inverted microscope body (IX83, Evident, Japan). The schematic setup is illustrated in Fig. 1. By incorporating a magnetic mirror mount, seamless switching between the two SRS lasers is possible. A 1.2 NA water-immersion objective lens (UPLSAPO60XW, Evident, Japan) is used to focus the pump and the Stokes beams into the sample. The back aperture is slightly under-illuminated (approx. 75%) in order to prevent any influence of differences in the beam diameters between the two laser sources on the effective numbers of Raman-active molecules in the focal spot. Light transmitted through the sample is collected by an oil-immersion condenser (U-AAC, NA = 1.4, Evident, Japan) and imaged onto a large photo-diode connected to a lock-in amplifier (dual frequency SRS detector set, 20 MHz detection band, MW Elektrooptik, Hamburg). Two short pass filters (F72-890, AHF anaylsentechnik, Tübingen) in front of the photo-diode ensure that only the pump light impinges on the detector and thus the stimulated Raman-loss is detected. The signal, demodulated by the lock-in amplifier is digitized with a National Instruments card (PCIe-7852, National Instruments, Austin) and visualized with the Imspector software package (Abberior Instruments, Germany). For rapid sequential imaging of various Raman bands (section 4.4), the oil-immersion condenser was replaced by a 0.55 NA air condensor (IX2-LWUCD, Evident, Japan).

**Fig. 1.**
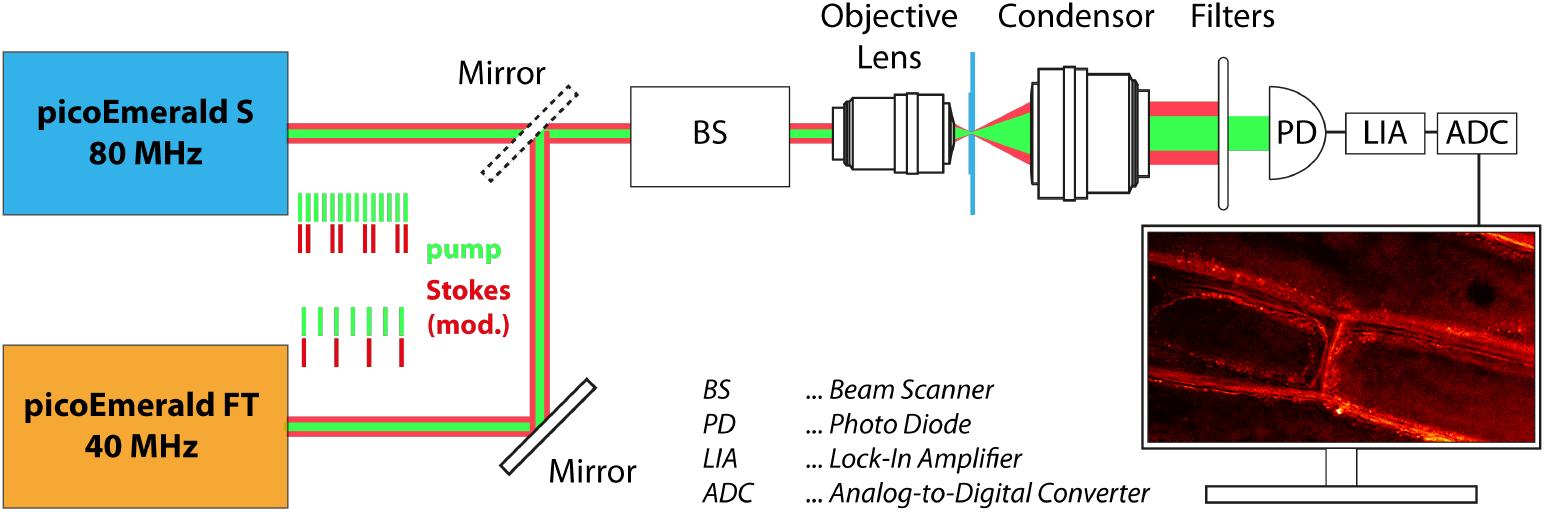
Schematic of the SRS setup: The two SRS light sources are directed through a beam scanner (BS) into an inverted microscope equipped with a water-immersion objective lens (NA 1.2). Transmitted light is collected by an oil-immersion condenser (NA 1.4) and analyzed by an SRS detection system consisting of a fast photodiode (PD) and a lock-in amplifier (LIA). A set of filters ensures that only the pump wavelength is detected. The LIA output is digitized using an analog-to-digital converter (ADC). A mirror on a magnetic mount enables seamless switching between the two light sources. The inset illustrates the pulse timing schemes for the pump and Stokes beams, highlighting the differences between the two laser systems.

### 3.3 Sample Preparation

All samples consist of a sandwich of two cover glasses of thickness No. 1.5, one with a size of 24 mm x 60 mm (0102242, Marienfeld, Lauda Königshofen) and one with 22 mm x 32 mm (631-0134, VWR International GmbH, Darmstadt), separated by imaging spacers (GBL654004, Merck, Darmstadt) that form a well with a diameter of 13 mm and a depth of 0.12 mm.

Rapeseed Oil sample: An imaging spacer (GBL654004, Merck, Darmstadt) is placed on the larger cover glass. Then, 20 µL of ordinary rapeseed oil is pipetted into the spacer’s well. The smaller cover glass is placed on top to seal the sample.

Onion Epidermal Cell sample: Epidermal cells from onions were gently extracted from between the scales and placed in the well. To ensure the epidermal layer remained as flat as possible for optimal imaging, the cells were placed onto a drop of deionized water within the well. The well was then gently filled with additional water, and the smaller cover glass was placed on top to seal the sample.

Oyster Prismatic Layer sample: To access the prismatic layer of oyster shells, the shells were carefully broken into pieces. Thin sections of the prismatic layer were selected and placed onto a drop of deionized water within the well. The well was then filled with additional water, and the smaller cover glass is placed on top to seal the sample.

## 4. Results

### 4.1 Laser source characterization

To accurately assess and compare the performance of both SRS lasers, it is essential to precisely characterize their noise behavior. Therefore, we performed laser intensity noise measurements following the procedures described by Audier et al. [8]. These measurements determine the laser’s relative intensity noise (RIN), which depends on both the average photodiode current and the frequency.

Figure 2 compares selected parameters of the picoEmerald S and picoEmerald FT lasers, shown in blue and orange, respectively. In figure 2a), the frequency-dependent RIN for both light sources, as well as the dark noise (gray), is presented. The measurements were performed with 10mW of laser light at 800nm impinging on a 50 Ω terminated photodiode (DET10A2, Thorlabs), leading to an average photocurrent, *I*_avg_, of 4.08 mA. The red line indicates the shot noise limit, calculated according to [8]:

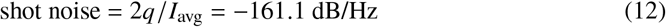

where *q* is the elementary charge. At a frequency of 20 MHz, both lasers operate in the shot noise-limited regime, which is where the modulation frequency of the Stokes beam is set.

**Fig. 2.**
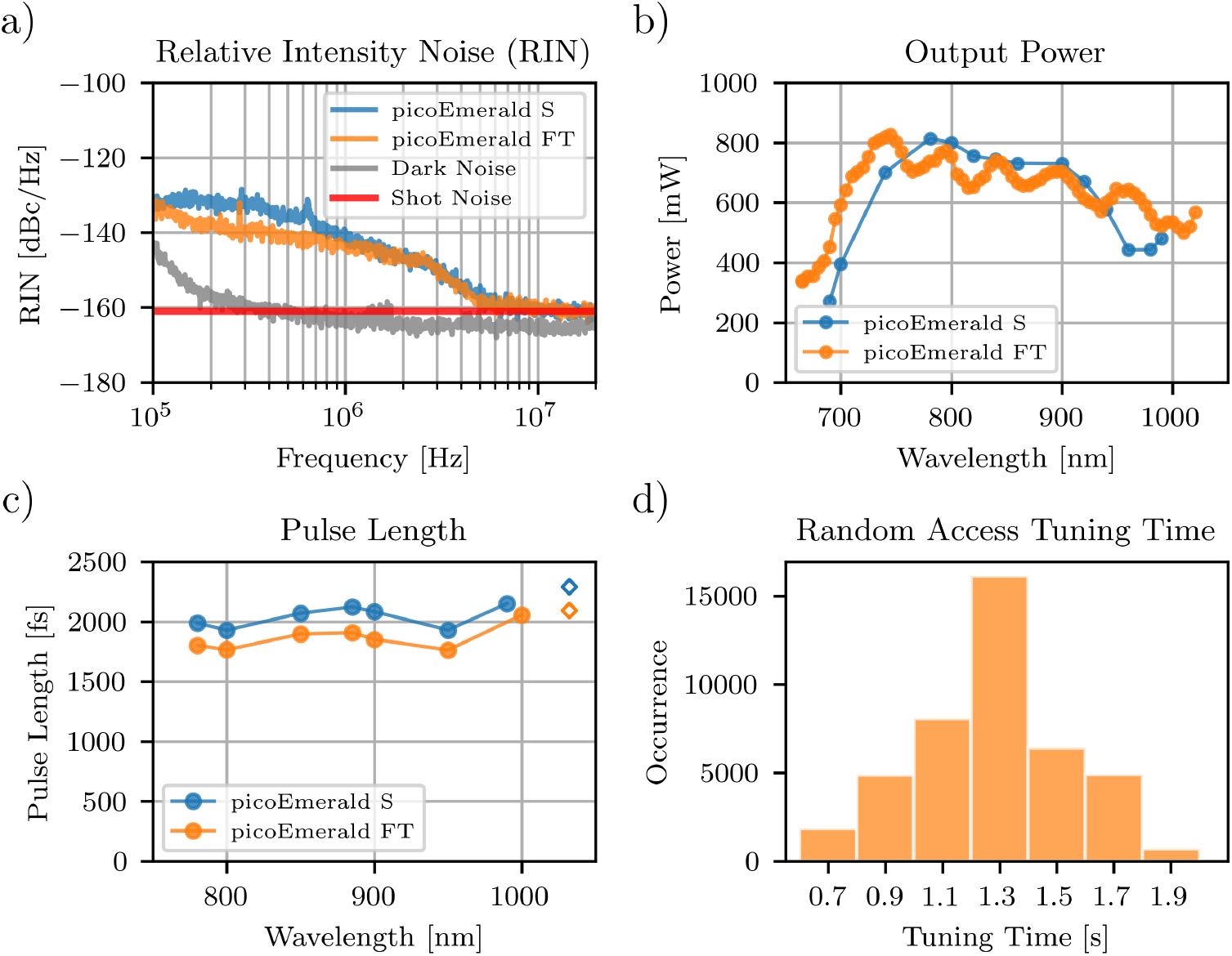
Comparison of performance metrics between the two SRS light sources: a) The newly developed laser (orange) demonstrates RIN levels comparable to the established system (blue). At the demodulation frequency of 20 MHz, both operate within the shot-noise-limited regime. b) Both lasers achieve output powers exceeding 500 mW across most of their tuning ranges, with the new system offering an extended tuning range. c) Both sources deliver pulse durations around 2 ps, with the new system achieving pulses that are, on average, 10% shorter. d) Performance tests with approximately 43,000 random wavelength accesses show an average tuning time of 1.3 s for the new laser system.

Figure 2 b) shows the optical output power of the picoEmerald FT over the entire tuning range (615nm to 1015nm). For the majority of the tuning range, the output power remains well above 500mW, which is sufficient to provide the tens of milliwatts of optical power required at the sample for SRS measurements [9, 12, 22].

The pulse durations of both SRS light sources across their tuning ranges, measured with an autocorrelator (Mini TPA/PD, APE, Germany), are shown in figure 2 c). Both sources maintain pulse durations of approximately 2 ps over the entire wavelength range, with the picoEmerald FT exhibiting slightly shorter pulses. The pulse durations of the Stokes beams are indicated by diamonds.

As described in sec. 3, the wavelength tuning mechanism of the picoEmerald FT was modified to allow for fast tuning times. For characterization, we tuned the OPO to random wavelengths within the entire tuning range approximately 43,000 times, and the time needed until stable operation was logged. A histogram of the tuning times (see figure 2 d)) shows that, on average, the laser stabilizes at any wavelength within the range of 660–1010 nm (210–5450 cm^−1^) in 1.3 seconds. A detailed comparison of both lasers is summarized in table 1.

**Table 1.** Detailed comparison of key parameters between the picoEmerald S and picoEmerald FT lasers.

|  | picoEmerald S | picoEmerald FT |
| --- | --- | --- |
| **Repetition Rate** | 80 MHz | 40 MHz |
| **Modulation Frequency** | 20 MHz<br>= 1/4 Repetition Rate | 20 MHz<br>= 1/2 Repetition Rate |
| **Bandwidth (pump & Stokes)** | 10 cm<sup>-1</sup> | 10 cm<sup>-1</sup> |
| **Tuning Range** | 700 nm - 990 nm<br>410 cm<sup>-1</sup> - 4600 cm<sup>-1</sup> | 660 nm - 1010 nm<br>210 cm<sup>-1</sup> - 5450 cm<sup>-1</sup> |
| **Output Power Stokes** | > 700 mW unmodulated<br>> 300 mW modulated | > 700 mW unmodulated<br>> 350 mW modulated |

### 4.2 Quantitative Comparison

To compare both laser systems quantitatively, we prepared a homogeneous rapeseed oil sample in accordance with section 3.3 and imaged consecutively with the picoEmerald FT and the picoEmerald S. Rapeseed oil contains a large number of *CH*_2_ and *CH*_3_ groups, which provide a strong stimulated Raman signal. The pump wavelength was tuned to 797nm to probe the *CH*_2_ symmetric stretch band at 2850 cm^−1^ [23]. We set the pump and Stokes powers at the sample position to comparable levels for both lasers; exact values are listed in table 2. The generated SRS signal was detected by a photodiode and isolated from noise using lock-in amplification with an integration time of 2 µs and a gain of 28 dB. We then digitized the signal at a sampling rate of 750 kHz. The offset voltage of the lock-in amplifier was adjusted such that the detected intensity was zero in the absence of incident light on the photodiode.

**Table 2.**
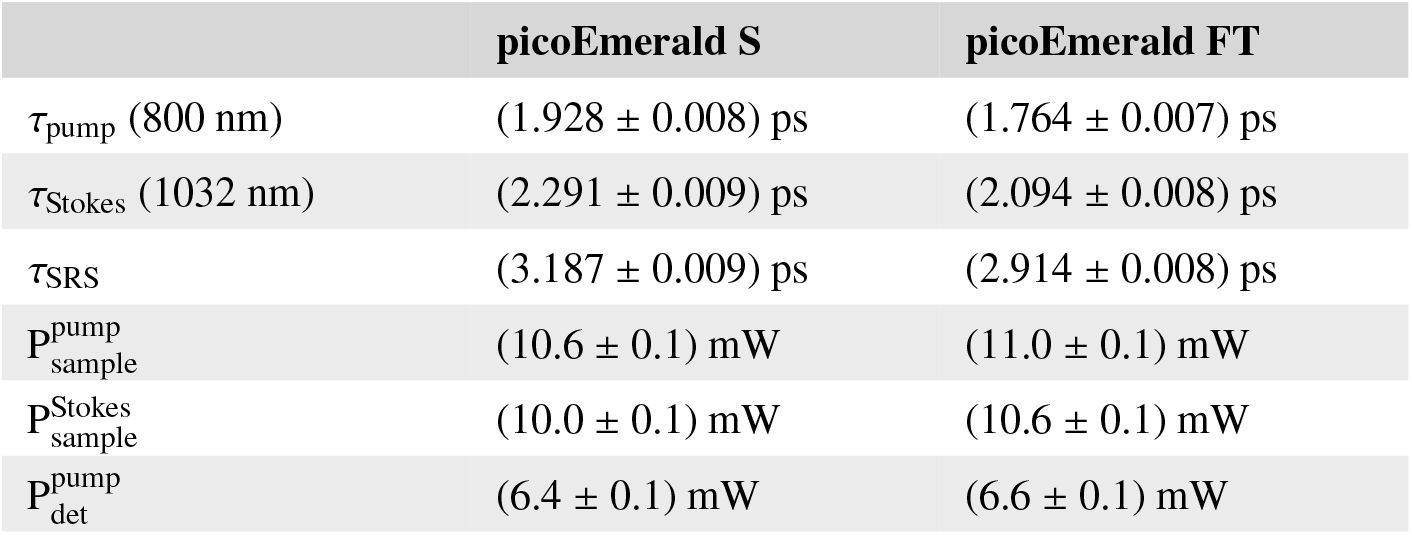
Laser powers at sample and detector position.

For both light sources, we acquired a series of data sets with varying pixel dwell times ranging from 2 *μ*s to 80 *μ*s. Each data set contained 25 consecutive measurements (frames) of the same field of view (40 *μ*m ×40 *μ*m), with a pixel size of 200 nm ×200 nm. To correct for signal fluctuations on the frame-to-frame time scale, the signal of each frame was normalized with respect to the sum of the first frame. For each data set, the average signal (*U*) and the variance of the signal (Var [*U*]) were calculated pixel-wise. By fitting histograms of *U* and Var [*U*] with Gaussian functions, their expected values and standard deviations were retrieved.

The data as a function of the pixel dwell time (τ_pdt_) for the measurements recorded by using the picoEmerald S and picoEmerald FT are shown in figure 3 in blue and orange respectively. In the top-left subplot, the average signals for using the picoEmerald S (*U*_S_) and the picoEmerald FT (*U*_FT_) are plotted. Data points represent the expected value, and error bars indicate the standard deviation, as determined from Gaussian fits. For both lasers the detected SRS signal was independent of the pixel dwell time. The arithmetic means and standard errors for the signals are: 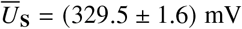 and 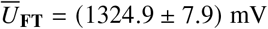.Calculating the ratio of 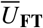 and 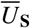 reveals that the signal in the measurements performed with the picoEmerald FT is increased by a factor of 4.02 ± 0.10. By using the measured pulse lengths and average powers in the focal plane (table 2) in conjunction with equation 9, we can calculate a theoretical ratio as follows:

**Fig. 3.**
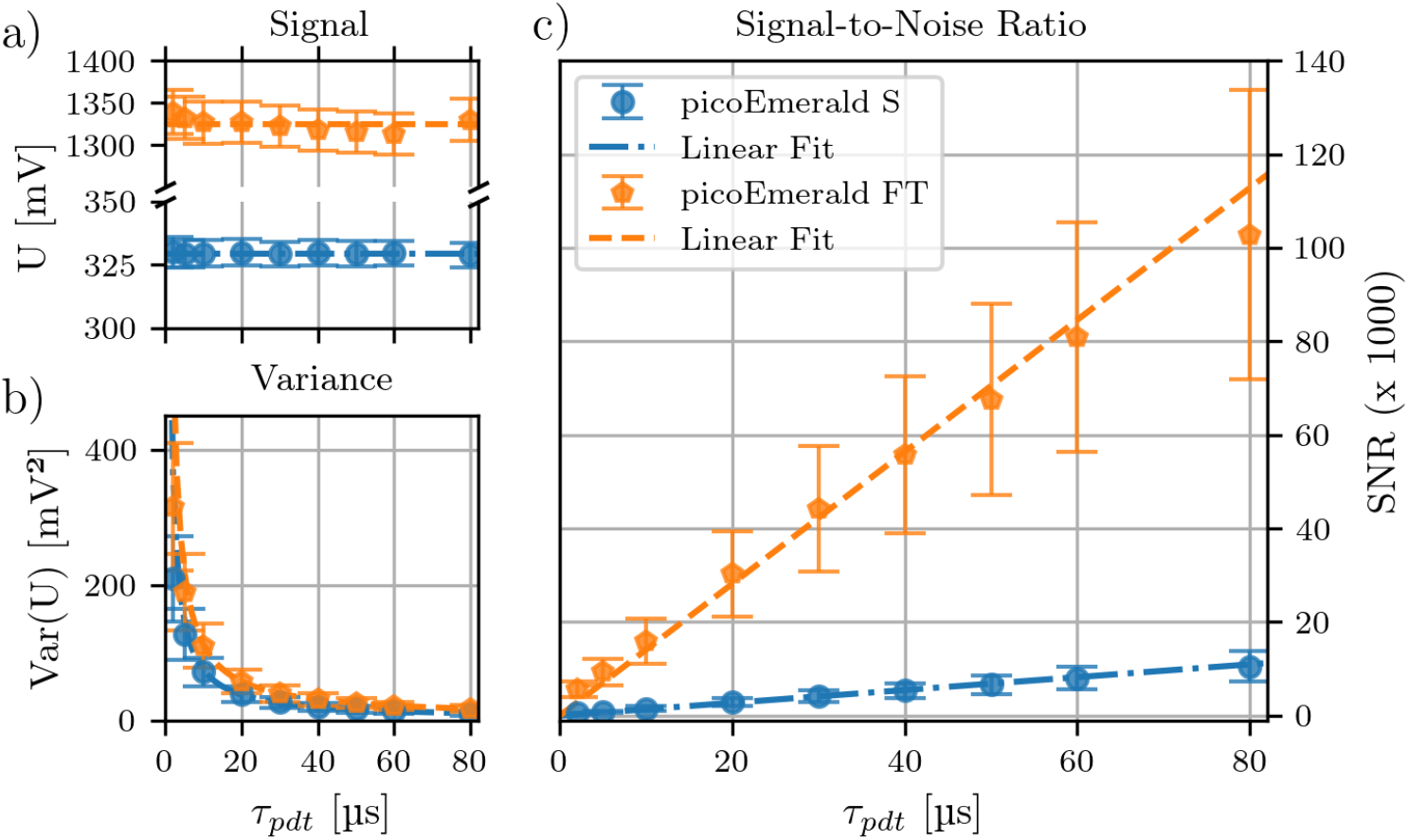
Quantitative comparison of signal, signal variance, and signal-to-noise ratio for the picoEmerald S (blue) and picoEmerald FT (orange) in SRS measurements of rapeseed oil: a) *U* shows a fourfold increase for the new laser. b) Var [*U*] is approximately 1.5 times higher for the new system. c) SNR increases approximately tenfold for the new laser, consistent with theoretical predictions. Data were acquired for varying pixel dwell times (2–80 µs).

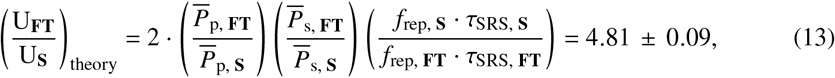

where the error is calculated by propagation of uncertainty of the individual parameters.

The bottom left subplot shows the variance of the signals (Var [*U*_*S*;*FT*_]). Both graphs are fitted with *f* τ_pdt_ = *a*/ τ_pdt_ +*b* and the result of the fits are shown as dotted lines. By calculating the ratio *a*_FT_ *a*_S_ it can be found that the signal variance of the picoEmerald FT is about 1.48 ±0.53 times larger than the signal variance of the picoEmerald S. Using the measurement parameters and equation 10, a theoretical ratio can be calculated for the shot noise limited case as:

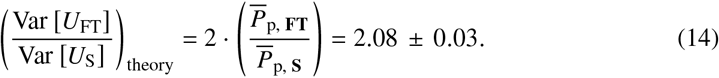

The graph on the right hand side shows the SNR calculated for the respective pixel dwell time according to SNR_SRS_ = *U*^2^ /Var [*U*] . As a proportionality of SNR_SRS_ ∝τ_pdt_ is expected, the data points are fitted with linear functions (dotted lines). To calculate the gain in SNR for the picoEmerald FT in comparison to the picoEmerald S, the ratio SNR_SRS, FT_ /SNR_SRS, S_ is calculated and shows a 10.29± 1.66 times increased SNR. Using equation 11, we can calculate the theoretically expected improvement factor as:

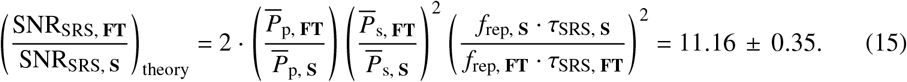

The results for the quantitative comparison of both lasers are summarized in table 3.

**Table 3.** Comparison of the picoEmerald S and picoEmerald FT.

|  | Experiment | Theory |
| --- | --- | --- |
| $U_{\text{FT}}/U_{\text{S}}$ | $4.02 \pm 0.1$ | $4.81 \pm 0.09$ |
| $\text{Var}[U_{\text{FT}}]/\text{Var}[U_{\text{S}}]$ | $1.48 \pm 0.53$ | $2.08 \pm 0.03$ |
| $\text{SNR}_{\text{SRS, FT}}/\text{SNR}_{\text{SRS, S}}$ | $10.29 \pm 1.66$ | $11.16 \pm 0.35$ |

### 4.3 SRS imaging of biological specimens

Under the previously described imaging conditions (see section 4.2), the picoEmerald FT demonstrated an approximately tenfold increase in SNR over the picoEmerald S. In order to investigate if the gain in SNR of the picoEmerald FT directly translates to an improved imaging capability in biological specimens, samples were prepared according to section 3.3, imaged, and analyzed qualitatively. We scanned a 120 µm x 120 µm field of view with a pixel size of 80 nm x 80 nm, with a pixel dwell time of 2 µs. The gain of the lock-in amplifier was set to 28 dB. In total, 128 consecutive images of the same field of view were acquired. By averaging *n*_average_ consecutive images, it is possible to generate an image with an effective pixel dwell time that is *n*_average_ times longer than the experimental pixel dwell time.

Figure 4 shows SRS images of the *CH*_2_ streching mode at 2871 cm^−1^ in epidermal onion cells. The pump wavelength was set to 796.1 nm and the laser powers at the sample position were 23 mW for the pump and 10 mW for the Stokes light. The upper row shows images acquired with the picoEmerald S and the lower row images acquired with the picoEmerald FT. In figures 4 a), c), e) & g) the full field of view is shown, while b), d), f) & h) show zoomed in regions, indicated by the dashed lines. While in the image acquired with the picoEmerald S at 2 µs pixel dwell time (fig. 4a and 4b) SRS signal around the cell walls is visible. Increasing the effective pixel dwell time tenfold (averaging over *n*_average_ = 10 consecutive images) reveals much greater details of the present sub-structure (c.f. fig. 4c and 4d). Remarkably, a similar level of detail is achieved with the picoEmerald FT at a 2 µs pixel dwell time (see fig. 4e and 4f) and increasing the effective pixel dwell time using the picoEmerald FT to 20 µs (*n*_average_ = 10, fig. 4g and 4h) increases the image quality further. Taken together, these images show that the picoEmerald FT achieves comparable image quality as the picoEmerald-S, but with one-tenth the acquisition time. With equal acquisition times, the SNR is boosted and even finer structures become discernable. This capability holds particular significance in SRS imaging, where molecules in the fingerprint region, known for their relatively low SRS signal, are frequently under scrutiny.

**Fig. 4.**
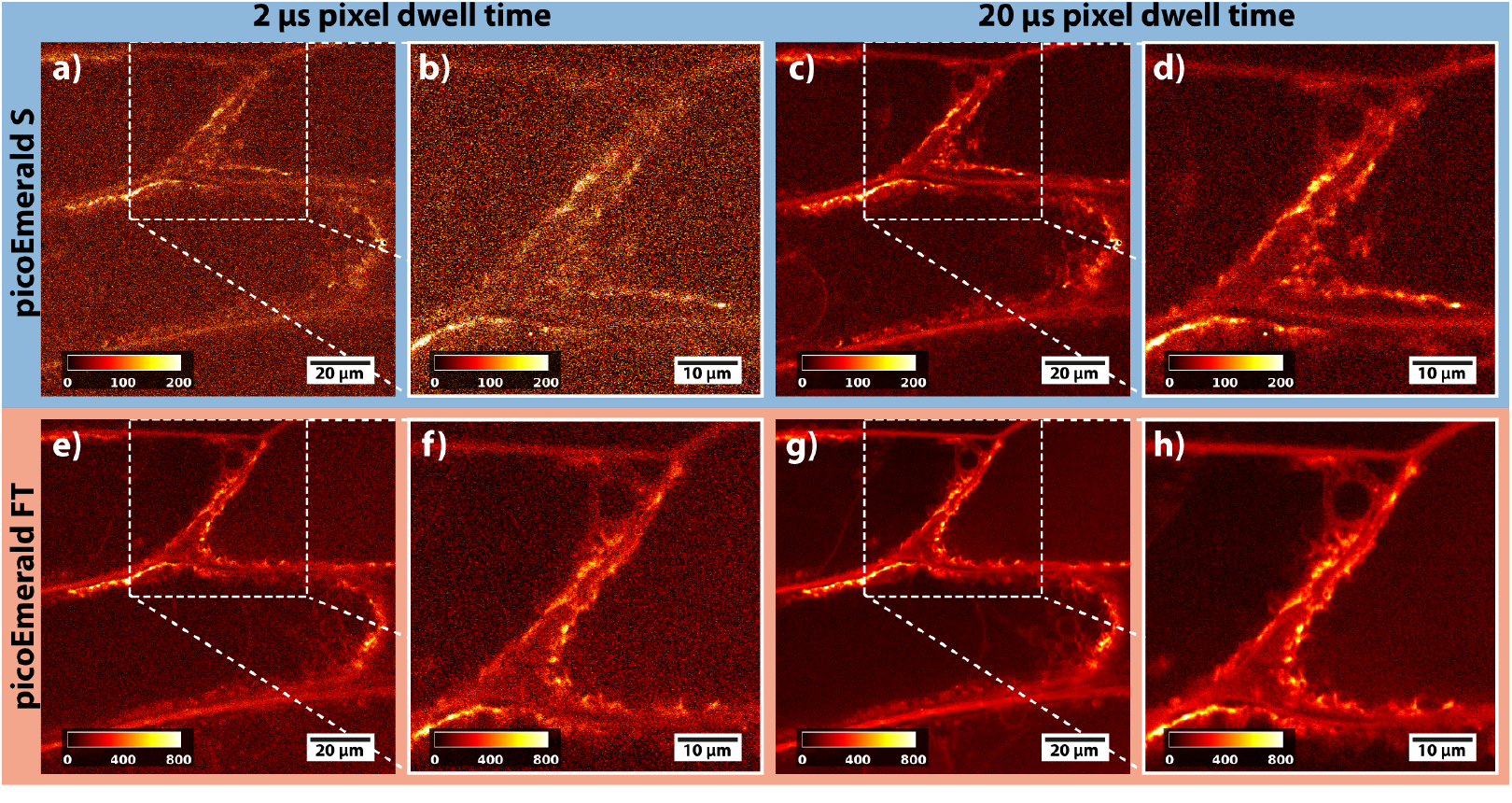
SRS images of the CH_2_ stretching vibration at 2871 cm^−1^ in epidermal onion cells. The upper row shows images acquired with the picoEmerald S, and the lower row shows those acquired with the picoEmerald FT. Each system includes an overview image and a magnified inset. Increasing the pixel dwell time from 2 µs (a, b, e, f) to 20 µs (c, d, g, h) significantly enhances image quality and feature visibility for both systems. Notably, at 2 µs dwell time, the picoEmerald FT delivers image quality comparable to the picoEmerald S at 20 µs dwell time. Further extending the pixel dwell time to 20 µs for the picoEmerald FT (g, h) results in even higher image quality.

To further explore the performance of the new laser source, we carried out a similar imaging experiment, this time using the amide I band in epidermal onion cells [11] (Figure 5). The pump wavelength was set to 880 nm to probe the vibrations at a 1670 cm^−1^ and the laser powers used are 30 mW and 9 mW for the pump and Stokes light respectively. In the 2 µs pixel dwell time images of the picoEmerald S (fig. 5a and 5b) the cell structures are faintly discernible, although the predominant aspect of the image is noise. As in the previous example, with a ten times increased pixel dwell time, the cells become visible (fig. 5c and 5d). The images acquired with the picoEmerald FT using a 2 µs pixel dwell time (fig. 5e and 5f) have a comparable image quality as the 20 µs picoEmerald S image (fig. 5c and 5d). By increasing the effective pixel dwell time to 20 µs for the picoEmerald FT (Figs. 5g and 5h) the highest image quality was obtained. A further example of imaging in the fingerprint region is shown in figure 6. Here, we measured of the crystalline calcite (CaCO_3_) band which is at around 1098 cm^−1^ [24]. The laser parameters were *λ* _*p*_ = 927.2 nm, *P*_*p*_ = 6 mW and *P*_*S*_ = 4 mW. As observed in the previous examples, images acquired with the picoEmerald FT exhibit a noticeable enhancement in both image noise and level of detail.

**Fig. 5.**
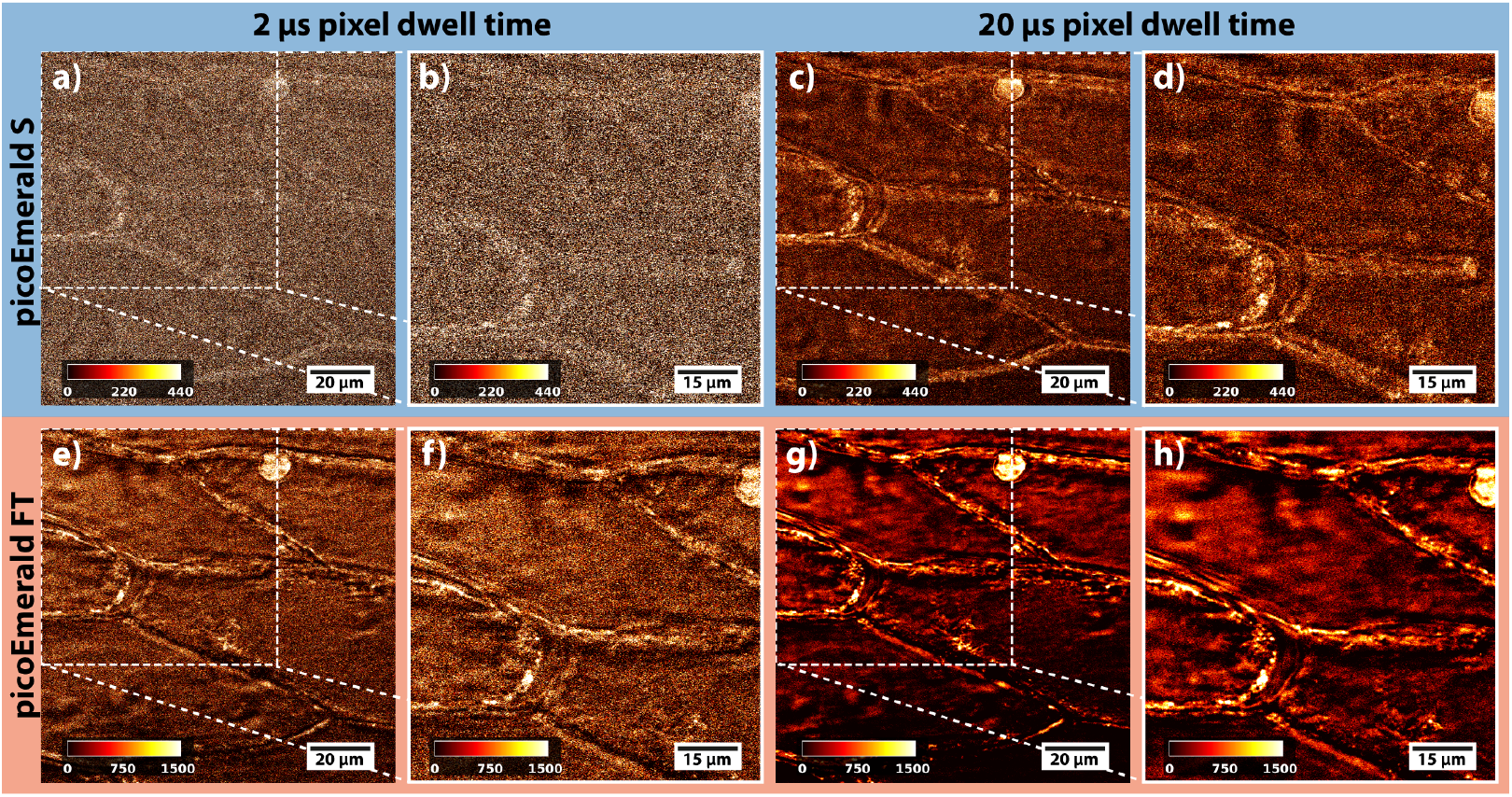
SRS images of the amide-I band at 1670 cm^−1^ in epidermal onion cells. The upper row shows images acquired with the picoEmerald S, and the lower row shows those acquired with the picoEmerald FT. Each system includes an overview image and a magnified inset. Increasing the pixel dwell time from 2 µs (a, b, e, f) to 20 µs (c, d, g, h) significantly enhances image quality and feature visibility for both systems. Notably, at 2 µs dwell time, the picoEmerald FT achieves image quality comparable to the picoEmerald S at 20 µs. Further increasing the dwell time to 20 µs for the picoEmerald FT (g, h) provides even greater detail.

**Fig. 6.**
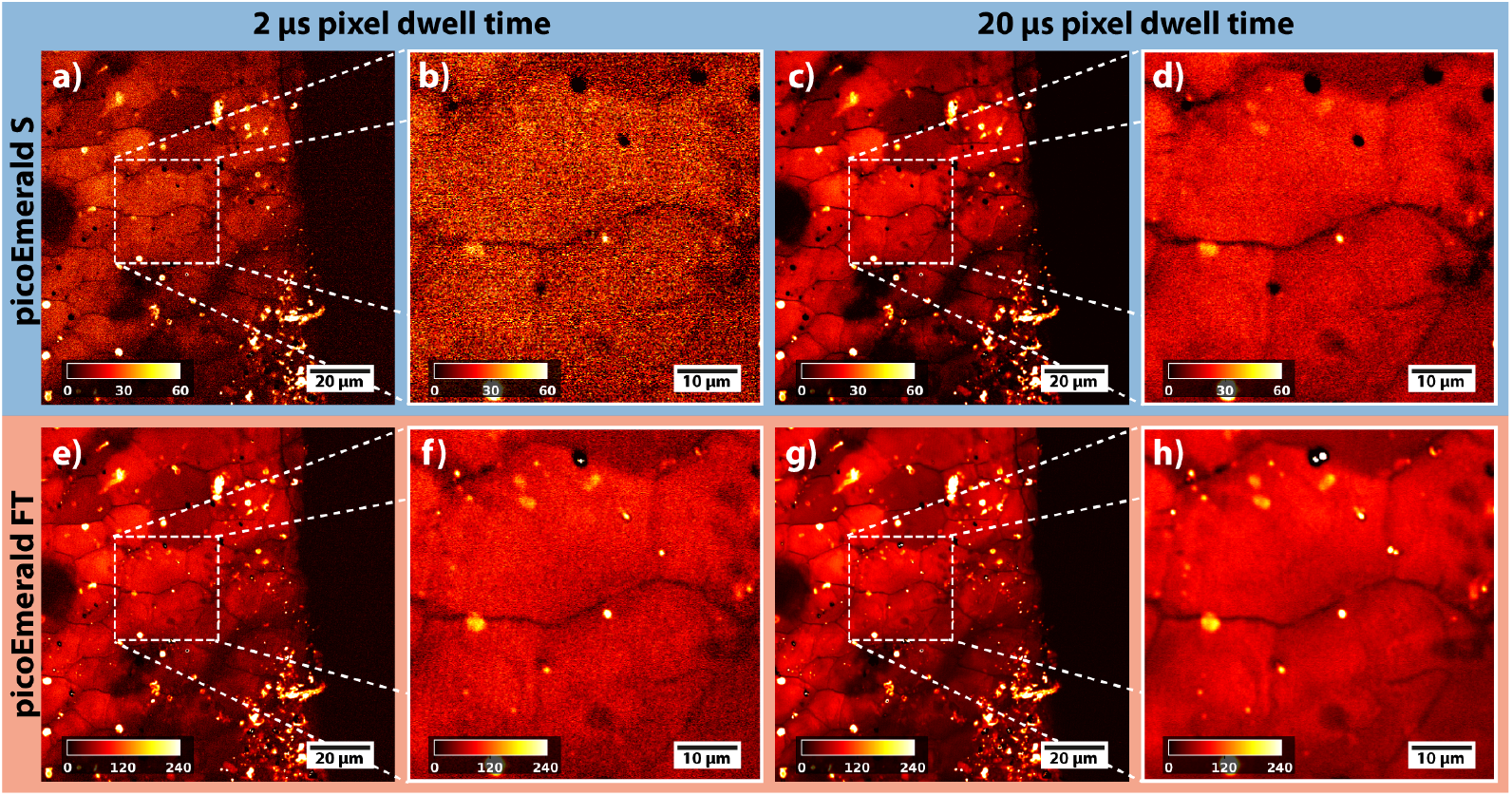
SRS images of the crystalline calcite (CaCO_3_) band at 1098 cm^−1^ in the prismatic layer of an oyster shell. The upper row shows images acquired with the picoEmerald S, and the lower row shows those acquired with the picoEmerald FT. Each system includes an overview image and a magnified inset. Increasing the pixel dwell time from 2 µs (a, b, e, f) to 20 µs (c, d, g, h) significantly enhances image quality and feature visibility for both systems. Extending the dwell time for the picoEmerald FT to 20 µs enhances the level of detail further.

### 4.4 Rapid sequential imaging of various Raman bands

With the development of the prototype picoEmerald FT, the OPO’s tuning mechanism was optimized, which sped up the tuning times from tens to hundreds of seconds down to an average tuning time of 1.3 s (c.f. section 4.1) and thus allowing for rapid sequential imaging.

For testing the tuning capabilities, we prepared samples containing epidermal onion cells (see section 3.3). We then sequentially tuned the pump wavelength to probe three Raman bands: the CH_2_ symmetric stretch (around 2870 cm^−1^), the CH_3_ stretch (at approximately 2930 cm^−1^), and the amide band (close to 1660 cm^−1^). The image acquisition was automated using a custom Python script.

For each image, we scanned a 100 µm × 100 µm field of view with a pixel size of 120 nm × 120 nm and a pixel dwell time of 2.8 µs. The gain of the lock-in amplifier was set to 48 dB. The laser powers in the focal plane were approximately *P*_*p*_ = 27.6 mW for all three pump wavelengths and *P*_*S*_ = 18.8 mW for the Stokes. The total measurement time was 16.6 s, exceeding the expected 9.6 s, which comprised approximately 1.9 s for imaging and 1.3 s for tuning per Raman band. This discrepancy arose primarily from overhead introduced by the Python scripting, including delays for shutter operations, acquisition initialization, and data storage.

Figure 7 shows SRS images of an onion cell captured at three distinct Raman bands, combined into a single RGB image to visualize the spatial distribution of different biochemical components. The RGB image is formed by assigning each Raman band to one of the primary color channels: red (*CH*_2_), green (*CH*_3_) and blue (amide) . Insets show the individual monochromatic images corresponding to each Raman band.

**Fig. 7.**
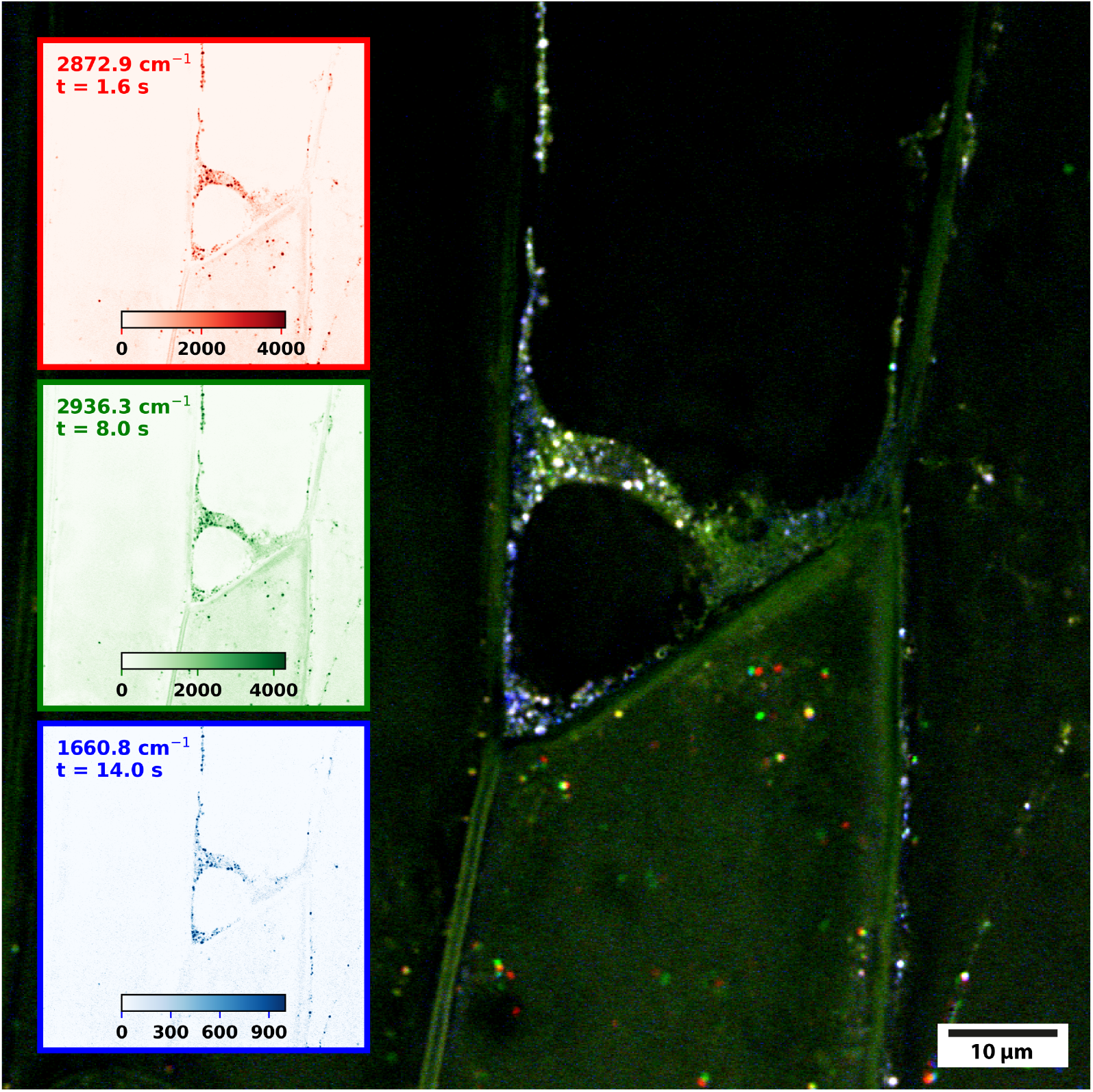
Rapid sequential imaging of Raman bands in epidermal onion cells. Insets depict individual SRS images of the CH_2_ band (2871 cm^−1^, red), CH_3_ band (2936 cm^−1^, green), and amide band (1661 cm^−1^, blue), annotated with the time elapsed since the start of the measurement. The composite image overlays the three bands to highlight the spatial distribution of the chemical components within the cells. The total measurement time for all bands was 16.6 s, showcasing the system’s rapid sequential imaging capabilities.

## 5. Discussion and conclusion

In this study we have evaluated the performance of two laser systems, the picoEmerald S and the prototype picoEmerald FT, for their applicability in stimulated Raman scattering (SRS) microscopy. By comparing these systems in terms of signal-to-noise ratio, tuning speed, and imaging performance, our work identifies technological advancements critical for high-resolution chemical and biological imaging.

The approximately tenfold increase in SNR achieved with the picoEmerald FT stems from its lower repetition rate, higher pulse energy, and optimized pulse duration. The strong alignment between measured SNR improvements and theoretical predictions validates the system modeling as well as the experimental approach.

The picoEmerald FT’s reduced tuning time of approximately 1.3 seconds represents a significant improvement, facilitating rapid sequential imaging across multiple Raman bands. This capability, demonstrated through the imaging of onion epidermal cells, allows the spatial mapping of multiple biochemical components in a single session, making it particularly valuable for analyzing heterogeneous biological samples.

In biological specimens, the picoEmerald FT resolved finer structural details within shorter acquisition times, particularly for weaker Raman signals in the fingerprint region. By achieving equivalent image quality with reduced exposure times at comparable laser intensities, the applied light dose is minimized, preserving sample integrity by reducing the likelihood of effects which could degrade sensitive specimens.

One remaining challenge is the overhead introduced by the current Python script-based control system, which delays wavelength tuning and image acquisition. These limitations, while manageable in a prototyping context, highlight the need for tighter integration into the microscope’s native control software. Such advancements would streamline the workflow and unlock the full potential of rapid sequential imaging for high-throughput studies.

In conclusion, the picoEmerald FT demonstrates clear advantages for SRS microscopy, particularly in applications requiring high sensitivity, rapid spectral imaging, and detailed structural resolution. Future work will focus on extending wavelength coverage and integrating advanced data processing tools, paving the way for broader adoption of SRS microscopy in biological and chemical research.

## Acknowledgments

The authors thank all partners of the NanoVIB consortium.

## Data availability

Data underlying the results presented in this paper are not publicly available at this time but may be obtained from the authors upon reasonable request.

